# Longitudinal analysis of influenza A virus deletion-containing viral genomes reveals key determinants of co-evolutionary dynamics and interference

**DOI:** 10.1101/2025.11.03.686333

**Authors:** Fadi G. Alnaji, Mireille Farjo, Shu Ling Goh, Daryl Zheng Hao AW, Paolo Alberto Lorenzini, Tongyu Liu, Rebekah Chan Lin-Zhen, Cornelius Lee, Marco Vignuzzi, Christopher B. Brooke

## Abstract

The substantial genetic diversity generated during influenza A virus replication facilitates both evasion of pre-existing host immunity and cross-species emergence. A major contributor to this diversity is the ubiquitous production of deletion-containing viral genomes (DelVGs) – viral RNAs with large internal deletions that arise during replication. DelVGs directly compete with wild-type (WT) genomes, and their accumulation has been implicated in modulating disease severity. However, the specific functional and genetic interactions between DelVGs and WT genomes remain poorly understood. To examine how DelVGs and WT genomes may co-evolve in mixed populations, we serially passaged influenza A virus under sustained high multiplicity-of-infection (MOI) conditions and used next-generation sequencing to build a longitudinal profile of DelVG emergence and dynamics. Early passages yielded a highly diverse repertoire of DelVGs across multiple segments, followed by a sharp contraction in overall diversity and the rise of one to two PB2-derived DelVGs that persisted at sustained high frequency. We identified a single-nucleotide substitution within PB2-derived DelVGs that significantly enhanced their replication and interference, whereas the same mutation proved lethal in the WT background. Collectively, these findings indicate that DelVGs are not mere byproducts of replication but dynamic genomic elements shaped by selection; their capacity to acquire adaptive mutations and outcompete WT genomes underscores their active role in shaping within-population viral dynamics.

## Intro

Influenza A viruses (IAV) are negative-stranded, segmented RNA viruses responsible for yearly seasonal epidemics and occasional global pandemics. IAV populations exist as highly diverse swarms of genetically and phenotypically heterogeneous particles (1). One of the most well-known contributors to viral population heterogeneity are defective interfering particles (DIPs): virions that carry one or more viral gene segments containing large internal deletions, also known as deletion-containing viral genomes (DelVGs) (2, 3). Because these deletions occur within essential viral genes, DelVGs cannot replicate independently, and depend on co-infection with WT virus to replicate.

IAV DelVGs are ubiquitous in both *in vitro* and *in vivo* infection models (4–8), as well as in clinical samples (9–12). Importantly, the abundance of DelVGs within clinical samples correlated negatively with infection severity in one study (12), emphasizing the need to better understand how DelVGs can modulate infection dynamics.

It has been known for decades that DelVGs can interfere with WT replication, leading to DelVGs outcompeting WT virus and numerically dominating viral populations under the right conditions (4, 13). The mechanisms by which this occurs are not fully understood, though competition with WT genomes for replication and packaging (14, 15) and stimulation of the innate immune response (16, 17) have all been proposed. We recently demonstrated that IAV DelVGs are actually replicated and packaged less efficiently than WT genomes over the course of a single replication cycle, raising questions about how they are able to outcompete WT virus over multiple generations (18).

To better understand the processes underlying DelVG accumulation and the evolutionary interplay between DelVGs and WT genomes within complex viral populations, we serially passaged IAV for 72 passages under sustained high MOI conditions and examined DelVG dynamics using next-generation sequencing (NGS). This approach yielded a detailed profile of the competition and co-evolution between DelVGs and WT genomes and revealed specific molecular features that govern the competitive success of DelVGs.

## Results

### Von Magnus oscillatory dynamics and DelVG distribution across the IAV genome are highly reproducible

To better understand how WT genomes and DelVGs interact and co-evolve within viral populations, we serially passaged recombinant A/Puerto Rico/8/1934 (rPR8) in two parallel replicate lineages under sustained high MOI conditions in MDCK cells. Both lineages were initiated from the same parental rPR8 population at an MOI of 10 TCID50/cell. Each passage was allowed to proceed for 18 hours, at which point half of the supernatant was used to infect the next passage while the other half was aliquoted and frozen.

To measure the accumulation of DIPs (packaged DelVGs) during passage, we measured infectivity by Tissue Culture Infectious Dose 50% (TCID50) assay, physical particles by hemagglutination (HA) assay, and calculated particle-to-infectivity ratios for each passage (**Fig 1A**). In both lineages, particle:infectivity ratios increased nearly five orders of magnitude over the first 7-9 passages, (compared with a ratio ∼10 for parental virus stock), suggesting a massive accumulation of DIPs relative to WT virus. Particle:infectivity ratios in both lineages crashed back to ∼10 over the next few passages, beginning an oscillatory pattern that continued throughout the 72-passage experiment. This oscillation pattern (often referred to as the “Von Magnus effect”) has been described many times before (4, 18–20), and our data indicates that this dynamic can be maintained stably within IAV populations for at least 100+ generations (assuming 2 viral generations per 18 hour passage).

**Fig 1.**
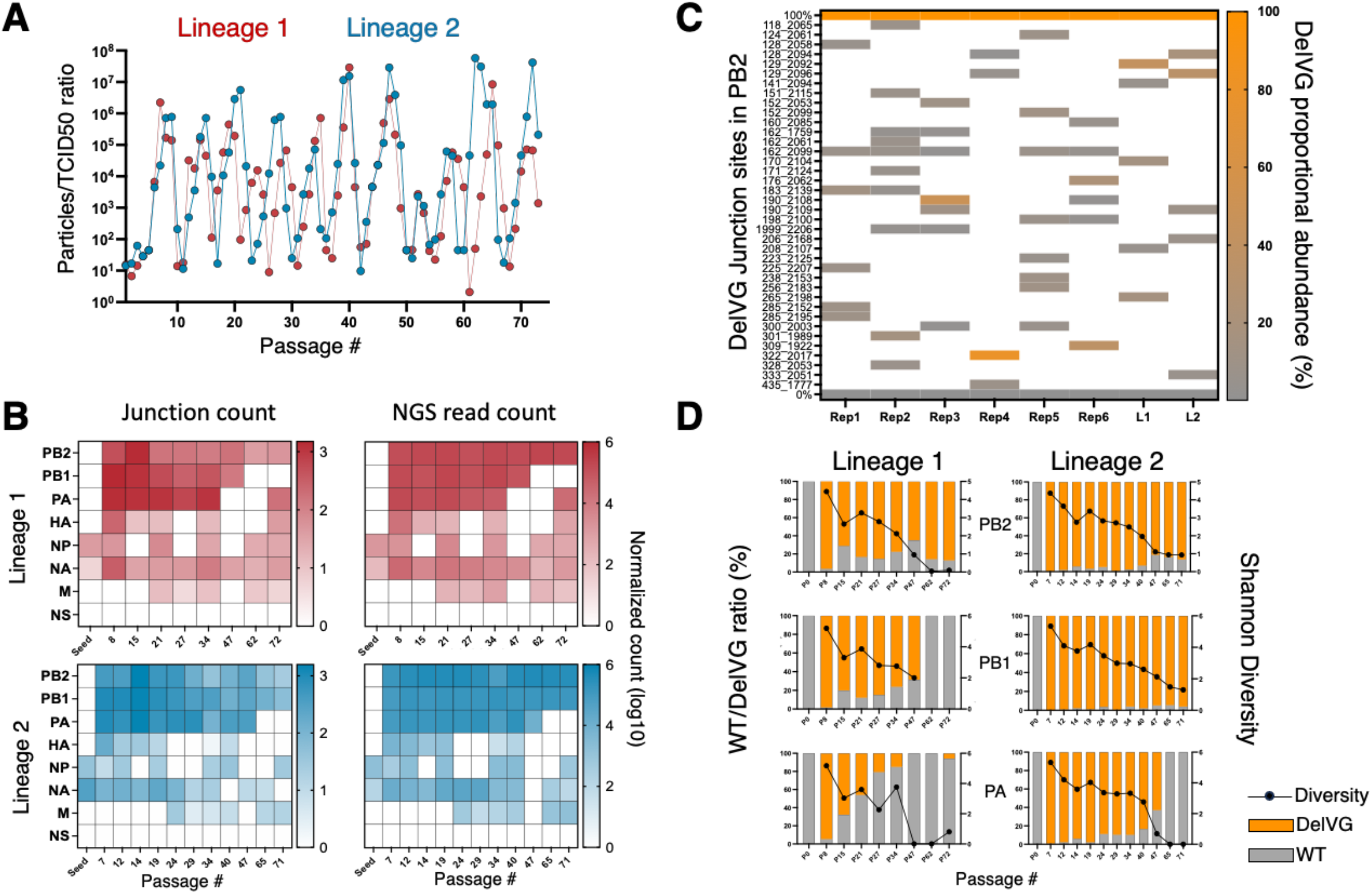
NGS-based frequency analysis of DelVG across all segments over time in various lineages. **(A)** Particles to infectivity ratios were calculated at each passage of both lineages (L1 & L2). The particles of each progeny were measured by the hemagglutination assay and divided by the infectivity, measured by TCID50. **(B)** Comparison of DelVG junction count (left panel) and DelVG NGS read frequencies (right panel) at each passage of all segments, both metrics were normalized to 106 mapped reads relative to the total aligned reads of each segment. **(C)** Heatmap showing the proportional abundance (%) of PB2-DelVGs, quantified from NGS reads, across six additional lineages together with L1 and L2. Only DelVGs reaching a relative abundance of ≥2% were analyzed. Each row represents a distinct junction and displays its abundance across the eight lineages. To facilitate interpretation, theoretical maximum and minimum values are shown at the top and bottom, respectively. White cells indicate absence of the junction in that replicate. **(D)** The WT/DelVG ratio was calculated (for PB2, PB1, PA segments) based on the NGS read counts and DelVG Shannon diversity was calculated and superimposed.

To identify the specific DelVGs that emerged within the passage populations, we used our optimized NGS-based pipeline (6) to characterize the deletion junctions in several representative passages with high particle:infectivity ratios (**Fig S1A**). Consistent with previous reports, hundreds of diverse DelVGs were detected across all segments except NS, with the majority derived from the polymerase segments (**Fig 1B**) (18–20). To further investigate the reproducibility of DelVG formation and accumulation, we initiated 6 additional replicate passage populations from the same parental population as lineages 1 and 2. We analysed the DelVG junctions in the polymerase PB2 segment present at passage 34 and observed minimal overlap in the specific DelVG deletions across the eight replicate lineages (**Fig 1C, Fig S1B**). This strongly suggests that while DelVG formation is reliably concentrated within well-defined hotspots and segments, the specific deletion junctions that form appear highly stochastic within those hotspots.

### DelVG diversity decreases over repeated passaging

We observed a reduction in both the numbers of unique deletion junctions (across all segments) and the numbers of DelVG-mapping NGS reads (in all segments except PB2 and PB1) in later passages (**Fig 1B**). To look at this another way, we plotted the WT:DelVG ratios for segments 1-3 (based on NGS read counts) across several passages where particle:infectivity ratios peaked, demonstrating that DelVG abundance dropped off after the initial expansion for some but not all segments (**Fig 1D**). This was not due to any variation in overall segment abundance, as WT versions of the three polymerase segments remained roughly equimolar throughout the experiment (**Fig S2A**).

While DelVG abundance remained high for some segments but not others, the diversity of individual junctions (based on Shannon entropy calculations) was sharply reduced for all segments (**Fig 1D**). Thus, even when DelVG abundance remained high for a given segment, it was due to a small number of specific deletions. Interestingly, the dramatic changes in DelVG segment distributions and genetic diversity that occurred over the course of the experiment did not disrupt the Von Magnus oscillation pattern, suggesting that the general pattern of WT-DelVG interactions was largely independent of the specific makeup of the DelVG population.

### DelVG competition dynamics differ between genome segments

To better understand the processes of competition and selection that shaped the DelVG populations during passages, we visualized the dynamics of individual DelVG junctions in relation to other DelVGs (**Fig 2**). DelVG dynamics differed substantially between genome segments but were generally similar between the two passage lineages.

**Fig 2.**
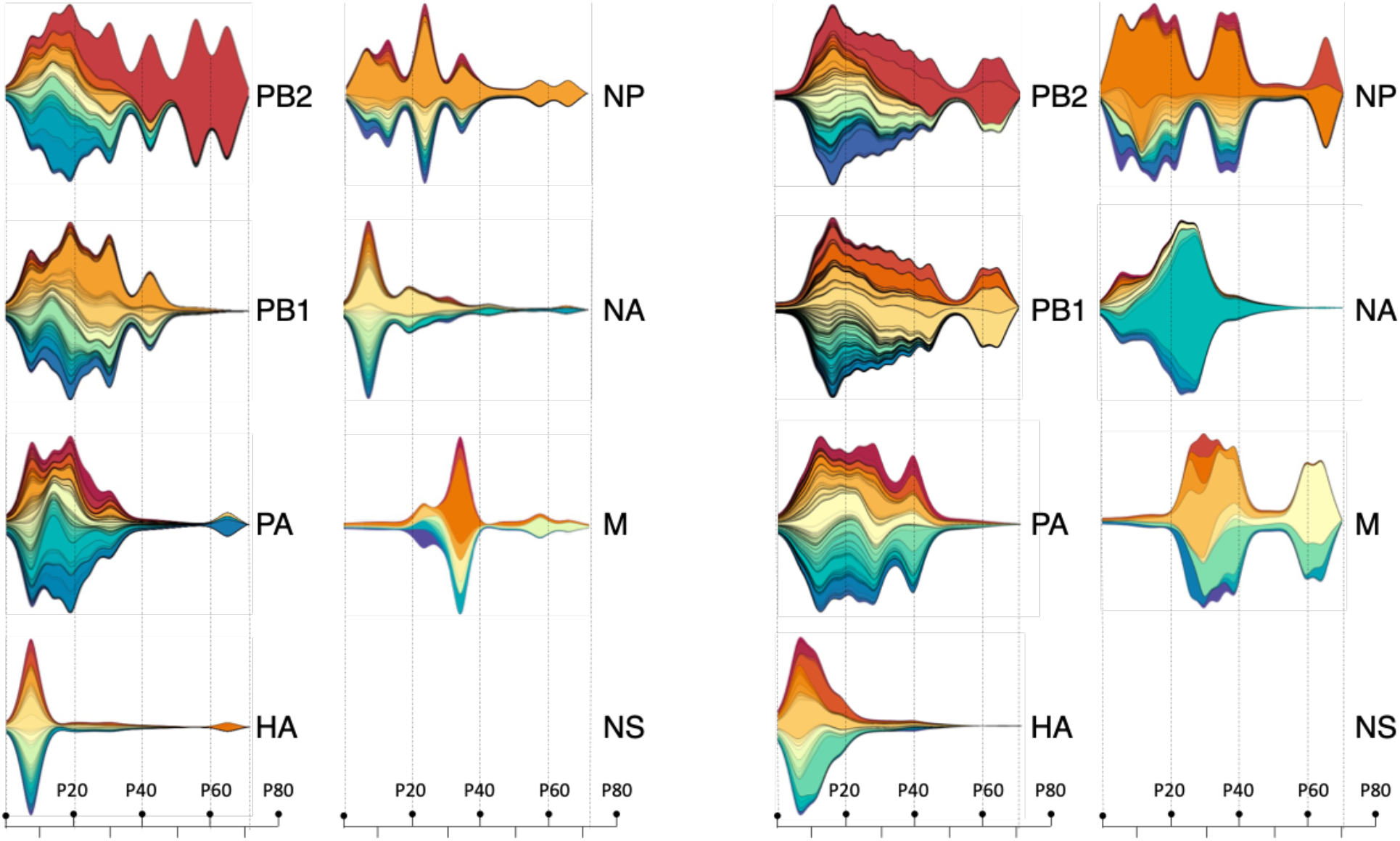
IAV PR8 DelVG evolutionary dynamics analysis over 72 passages in two lineages. Streamplots were generated with ggplot2 to depict DelVG competition dynamics. Each colored area corresponds to a unique DelVG junction; its area size at a passage reflects that junction’s proportional abundance (NGS reads for the junction/total DelVG-derived reads of the segment at that passage). At each passage, the vertical extent of the full stack is proportional to the segment’s total DelVG reads at that passage (height = K × total reads), with a single K used within each segment, determined by ggplot’s autoscaling based on that segment’s maximum total reads. Colors are assigned independently per plot; identical colors across segments do not denote the same DelVG.

For the PB2 segment, we observed the expansion of a diverse population of hundreds of distinct DelVGs in both lineages over the first ∼20 passages, followed by 1-2 specific DelVGs slowly and steadily taking over the population by the last passage (**Fig 2**). Notably, the three PB2 DelVG junctions that collectively took over lineages 1 and 2 had highly similar junction coordinates (129_2092, 128_2094, and 129_2096) (**Fig S2B**), suggesting that specific junction attributes may be associated with competitive success. The PB1 and PA segments showed similar initial expansions of diversity; however, the frequencies of individual deletions relative to each other appeared more stable, with no individual junctions outcompeting the others. Unlike PB2, DelVGs derived from PB1 (in lineage 1) and PA (in both lineages) largely disappeared after passage ∼50. HA DelVGs exhibited a rapid expansion of diversity, like the polymerase segments, but also exhibited a much more rapid collapse, largely disappearing from lineage 1 by passage 20 and lineage 2 by passage 40. NA DelVGs similarly peaked early and became largely absent after passage 40. Dynamics in the M segment were unique, in that DelVGs did not begin to accumulate until much later (first peaks around passage 20). The expansion and contraction dynamics of individual M-derived DelVGs also appeared to be more stochastic than those observed for other segments.

Altogether, our data suggest that DelVGs from different segments are subject to distinct evolutionary dynamics and that evidence of strong selection/competition dynamics is only clearly apparent in the PB2 segment. The near-total disappearance of DelVGs from some segments and lack of emergence of new DelVG junctions in later passages also suggest that viral populations can become resistant to invasion by *de novo*-generated DelVGs.

### Molecular characteristics of DelVGs over time

DelVG length has been suggested to govern competitive success, with shorter DelVGs thought to outcompete longer DelVGs and WT genomes (22). Similarly, the specific translation products of DelVGs have also been suggested to mediate interference with WT replication (23). We thus asked whether deletion size or post-deletion translation reading frame correlated with DelVG competitive success in our dataset. We compared DelVG sizes for segments 1-3 across four DIP-rich passages from both lineages (P8, P21, P34, P72) and found no significant difference in sizes (ANOVA test, P=0.2-0.4) (**Fig S3A**). We also found no evidence for enrichment of specific translation frames (**Fig S3B**). Although there was a notable bias toward the +0 frame at passage 72 in L1 (P < 0.001, Chi square), this was not seen in lineage 2 and may simply be an artifact of the small number of DelVG junctions remaining by passage 72. These observations suggest that under our experimental conditions, DelVG competitive dynamics cannot be simply explained by deletion size or post-deletion translation reading frame.

### Reconstitution of DelVG competition dynamics

It was possible that the complex competition and evolutionary dynamics we observed in our passage experiment were largely stochastic and were thus not reproducible. To test the reproducibility of the DelVG dynamics we observed, we attempted to recapitulate the competition dynamics we observed in the original experiment using recombinant, clonal DIP populations.

We selected three DelVGs from passage lineage 1, based on their competitive dynamics over time: DI250, which outcompeted all other DelVGs; DI143, which was less competitive but still maintained at detectable frequencies through most of the experiment; and DI289, which was rapidly outcompeted and disappeared (**Fig 3A**). We generated clonal recombinant DIPs encoding these specific DelVGs using reverse genetics, and developed qPCR primer probe sets that could distinguish between the individual DelVGs (**Fig S4**). We then performed competition experiments in which MDCK cells were co-infected with each of the three DIPs at an MOI of 1 PB2 genome equivalent (PB2-GE) per cell (for a combined total of 3 PB2-GE/cell), together with WT PR8 at an MOI of 10 PB2-GE/cell, in three independent lineages. Control infections lacking WT virus confirmed that the recombinant DIPs did not replicate, as expected. (**Fig S4**). We performed 16 undiluted passages of these populations and quantified the absolute abundances of DI250, DI143, DI289, and WT PB2 segment at each passage by RT-qPCR (**Fig 3B**).

**Fig 3.**
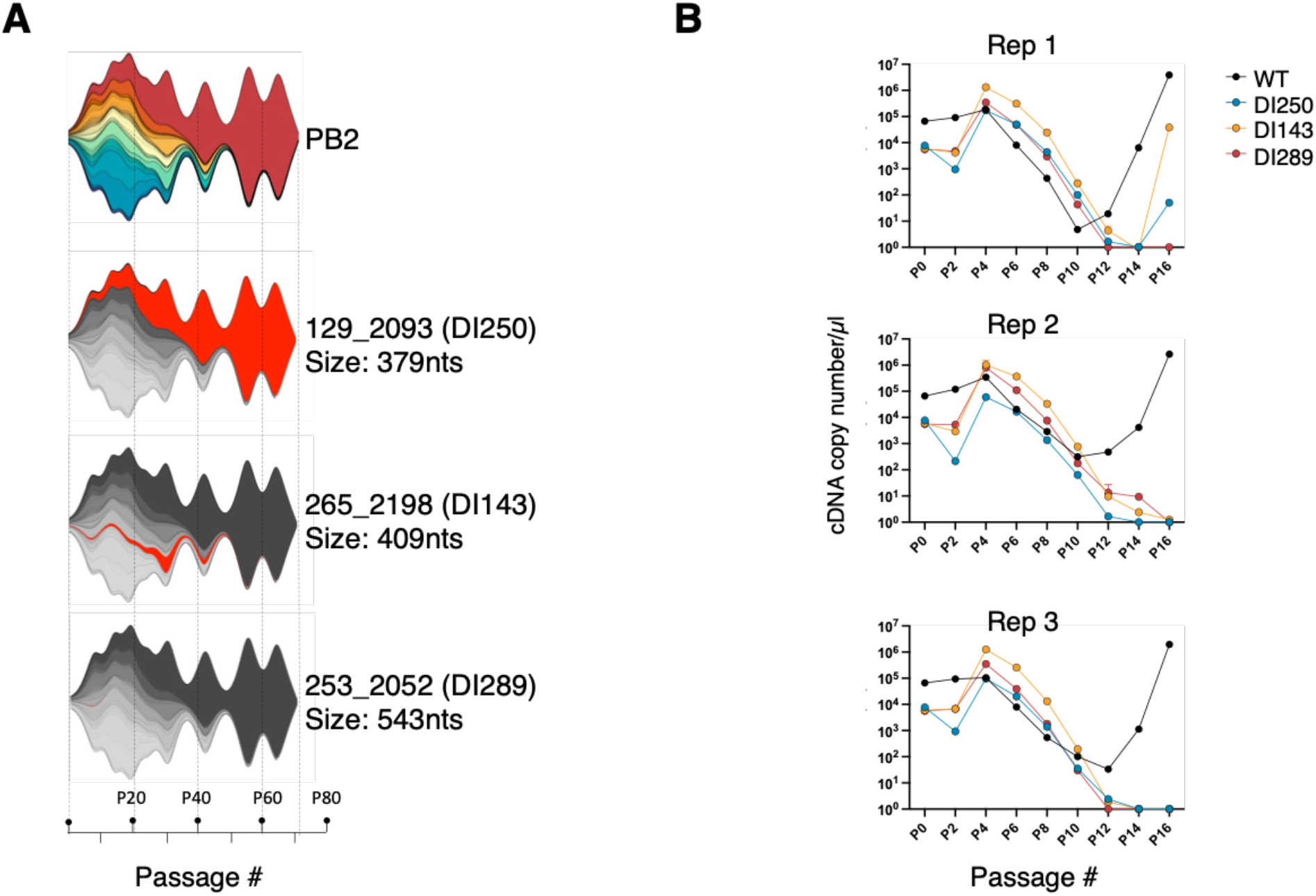
Competitive dynamics of clonal DelVGs during serial passage. **(A)** The top streamplot is the original L1-PB2 evolutionary dynamics analysis and was placed here for reference. The remaining 3 streamplots highlight the DelVG we picked in red with their names, junction sites, and sizes placed next to them. **(B)** MDCK cells were co-infected with clonal DIPs (DI250, DI289, DI290) at an MOI of 1 each based on PB2 gene equivalents (total MOI of 3), together with WT virus at 10 MOI, and serially passaged for 16 passages.

In broad strokes, these competition experiments reproduced the expected WT-DelVG dynamics, where the DelVGs outcompete the WT virus over the first several passages, before being overtaken by WT virus during the final passages. When we examined the dynamics of individual DelVGs, however, these conditions failed to recapitulate the specific inter-DelVG competition dynamics observed in the original experiment. While DI250 dominated in the original passage experiment, it was consistently outreplicated by DI143 and DI289 in all replicate experiments. Across the three replicates, DI143 consistently replicated to higher titers than the other DelVGs. These data demonstrated that DelVG competition dynamics can be highly reproducible across independent experiments, at least under artificial, reductionist conditions, suggesting that individual DelVGs have intrinsic competition phenotypes that can drive reproducible dynamics. However, these results also indicated that our competition experiments failed to recapitulate key aspects of the original passage experiments that gave rise to the dynamics we observed.

### Emergence of DelVG-associated SNPs during WT/DelVG co-evolution

Our inability to reproduce the DelVG competition hierarchy observed in the original experiment using recombinant DIPs suggested that we were failing to reproduce a key driver of the original dynamics. We hypothesized that competition between WT and DelVGs may have driven the emergence of substitutions that influenced WT/DelVG dynamics. To test this, we developed a bioinformatics approach to distinguish the reads of the selected DelVGs from other DelVGs and WT in the sequencing data generated from the long passage experiment, aiming to identify single nucleotide polymorphisms (SNPs) that emerged within DelVGs over the course of passaging (**Fig 4A**).

**Fig 4.**
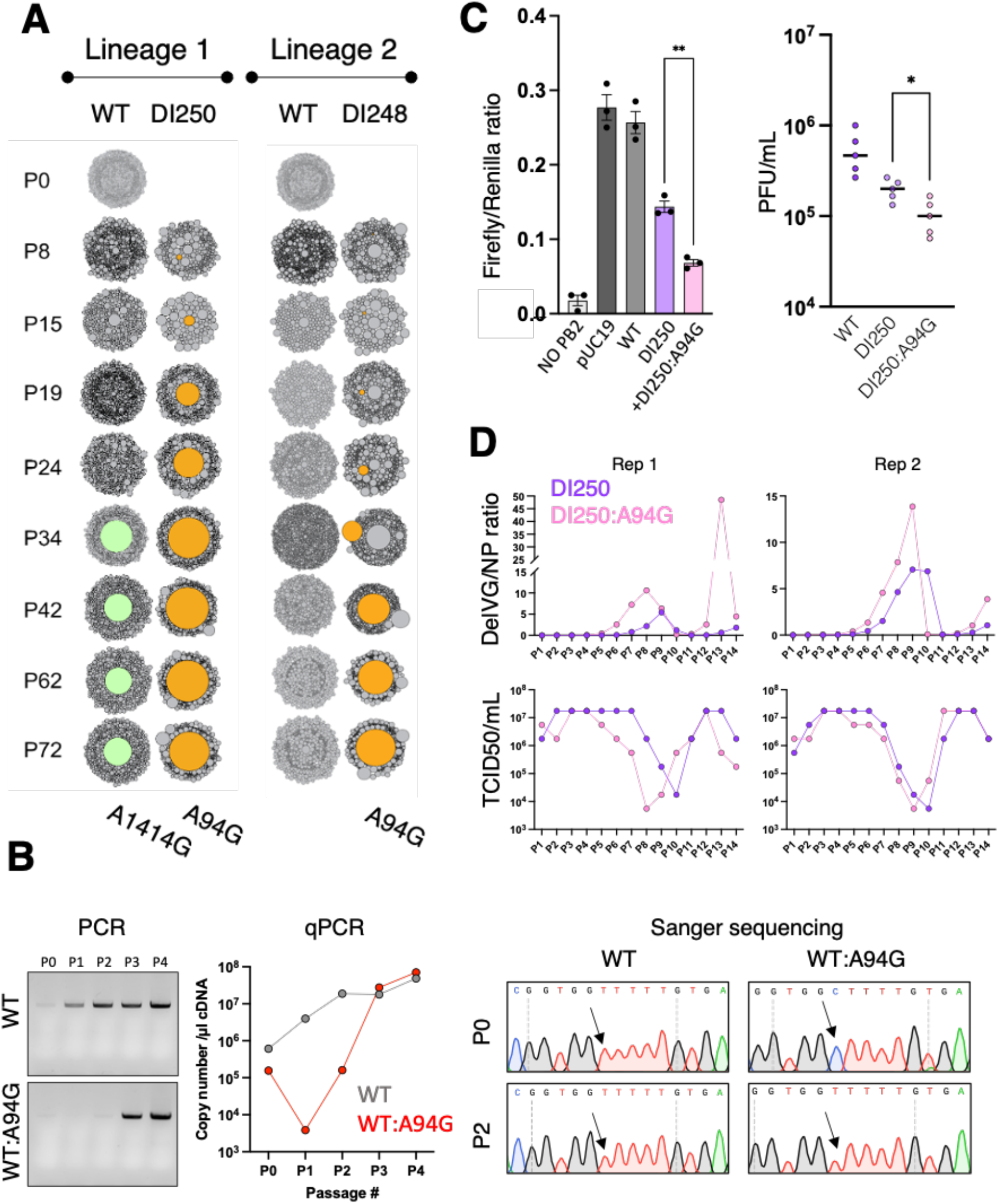
The effect of the SNP A94G in WT and DelVG. **(A)** A circle-packing diagram visualizes the mutation frequencies in both the WT virus and the major two dominating DelVGs in both lineages. Each circle represents a distinct SNP, with its size proportional to the mutation frequency. The most frequent mutations are listed below the figure and highlighted as light green (WT) and orange (DelVG). **(B)** The impact of the A94G mutation on the WT virus was illustrated through agarose gel, qPCR & Sanger sequencing using PB2-specific primers. **(C)** Left panel, the minigenome replicon assay is employed to measure the polymerase activity in the presence of DI250 or DI250:A94G. pUC19 was used as a negative control. Right panel, competition assay was performed between WT virus and clonal populations of either DI250 or DI250:A94G, using a 1:10 WT:DIP input ratio normalized by NP copy number. Progeny from these infections were subsequently quantified by plaque assay. **(D)** 293 cells were transfected with 500ng plasmids that express both the negative and positive strands of DI250 or DI250:A94G. Following the transfection, the virus was introduced 24 hours later. After another 24 hours post-infection, 50µl of the viral progeny was used to initiate infection in fresh MDCK cells, a process repeated 14 times. The proportion of DelVG to NP segment was determined using qPCR, shown in the top panel, while the TCID50 for each passage was assessed and displayed in the bottom panel. The experiment was carried out in duplicate. Statistical significance was determined by Welch’s t-test (*, p = 0.01; **, p = 0.001).

In lineage 1, a single SNP rose to high frequency within the dominant DelVG DI250 after passage 15 (**Fig 4A**), changing adenine to guanine at position 94 of the (+) strand sequence (hereafter referred to as A94G), which coded a threonine-to-alanine substitution at position 23 of PB2. This same SNP also arose to high frequency within one of the dominant DelVGs in lineage 2, DI248, suggesting that this SNP may confer some advantage during WT/DelVG co-evolution. Interestingly, this SNP was absent from WT PB2-derived reads. When introduced into WT PR8 by reverse genetics, the A94G substitution markedly attenuated viral replication, and rapidly reverted to WT (**Fig 4B**). These findings suggest that while A94G may be tolerated – or even advantageous – within DelVG backgrounds, it is maladaptive in WT viruses, underscoring its context-dependent effect on viral fitness. Consistent with this, A94G nevertheless rose to high frequency in competitive DelVGs in both passage lineages despite being highly attenuating in WT PR8 (**Fig 4A**).

### A94G enhances DelVG replication kinetics and inhibitory activity

To better understand the effects of the A94G mutation on DelVG function, we measured the relative abilities of DI250 and DI250:A94G to interfere with viral polymerase activity in a minigenome replicon assay (**Fig 4C, left panel**). In this assay, we expressed the components of the viral polymerase complex from plasmids in cells, along with a firefly luciferase-encoding reporter RNA derived from the viral NS gene segment and measured the accumulation of firefly luciferase signal as a surrogate for viral polymerase activity. We found that DI250 inhibited polymerase activity, relative to a WT PB2 control, and that this inhibitory effect was significantly increased for DI250:A94G.

To assess whether the outcomes observed in our minigenome assay were recapitulated during viral infection, we re-generated DI250 and newly generated DI250:A94G, producing DIP clonal populations of both (**Fig S5A**). Next, we used these DIPs to conduct a competition assay at a WT:DIP ratio of 1:10 NP-GE/cell (**Fig 4C, right panel**). Our finding mirrors the minigenome data, offering further support that A94G enhances the interfering capacity of DI250.

Finally, to directly examine the effects of A94G on DelVG competition dynamics, we co-passaged WT PR8 along with either DI250 or DI250:A94G for 14 passages and measured both DelVG abundance (by RT-qPCR) and fully infectious particle yield (by TCID50 assay). The relative abundance of DI250:A94G increased earlier and rose to a higher magnitude compared with DI250 (**Fig 4D**). Consistent with this, we observed that fully infectious virus titers declined more rapidly in the presence of DI250:A94G compared to DI250. Importantly, Sanger sequencing at late passages confirmed the persistence of A94G (**Fig S5B**), indicating that it is stable within DelVGs. This suggests that the emergence of this substitution enhances the competitive advantage of DI250 and likely facilitated the evolutionary success of the DI250 genotype during long-term passaging.

### A94G has no clear effect on packaging

To better assess the effects of A94G on DelVG replication and packaging, we performed an independent passaging experiment in which WT PR8 was co-passaged with either clonal DI250 or DI250:A94G individually, using an input of 0.1 MOI for each DIP and 1 MOI WT (PB2-GE/cell) in MDCK cells, with duplicate lineages for each condition. Quantitative tracking began at P3, when both DelVGs were reliably detectable at comparable levels across lineages, providing a common baseline (**Fig 5A**). From P3 onward, DelVG levels were measured in cells (6 hpi) and in virions up to P5 by junction-specific qPCR, with NP assayed in parallel as a reference. This allowed us to independently examine replication and packaging, enabling direct comparison of the two DelVGs under matched conditions. Across all time points and replicates, DI250:A94G exceeded DI250 in both intracellular and virion-associated copy number (at comparable NP), indicating higher replication efficiency.

**Fig 5.**
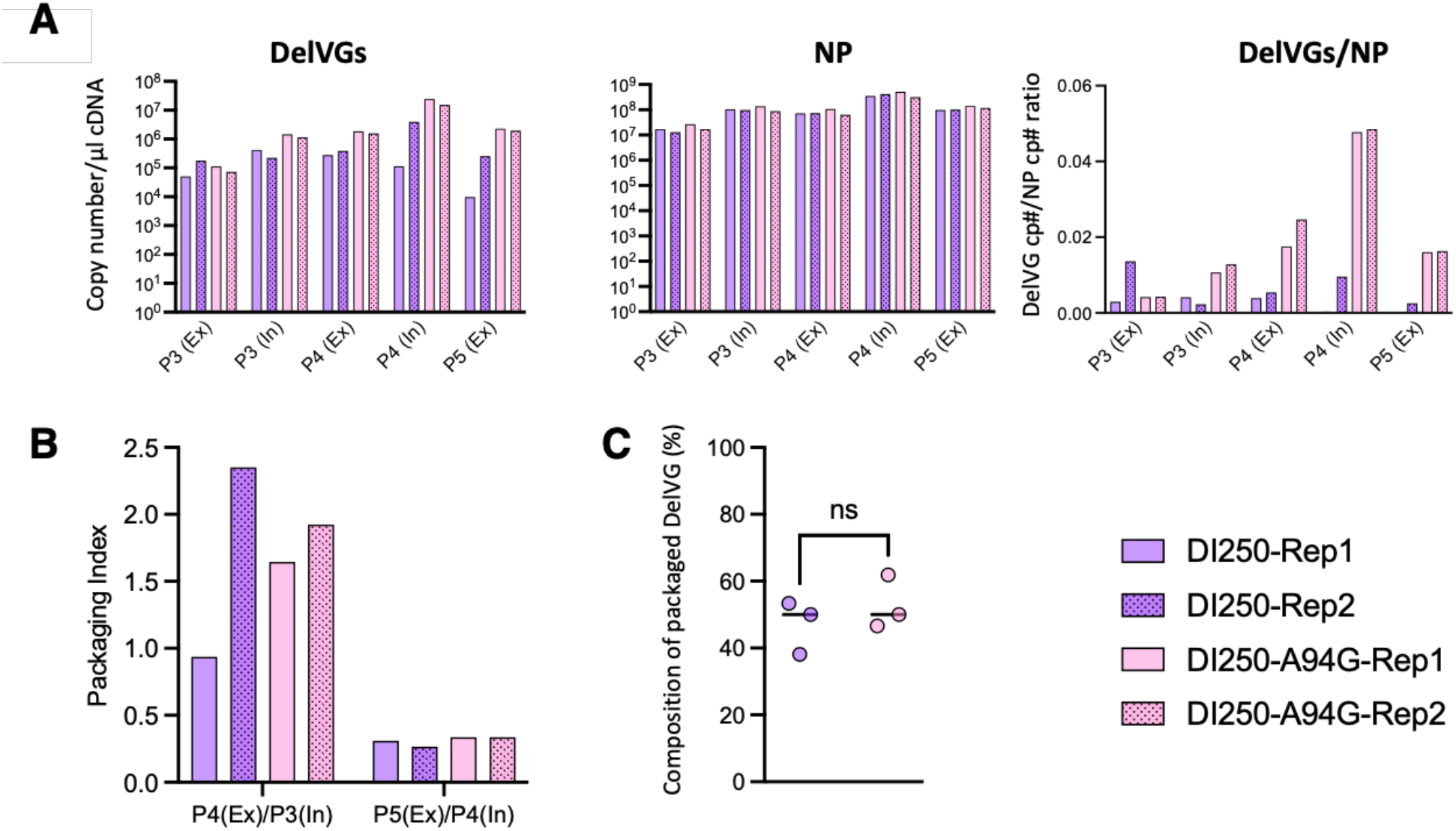
DI250 and DI250:A94G replication and packaging dynamics. Clonal DI250 or DI250:A94G populations were co-passaged with WT PR8 for five serial passages each in duplicate lineages, individually. **(A)** The left panel displays DelVG copy numbers, the middle panel shows NP vRNA copy numbers for each condition, and the right panel presents the DelVG-to-NP ratio. By passage 3 (P3), DelVG levels were comparable across samples. From this point onward, intracellular (In, 6 hpi) and extracellular (Ex, 24 hpi) dynamics were tracked through passage 5 (P5). Replicate 1 is shown in solid color and replicate 2 in the same color with a patterned fill. **(B)** Packaging index was calculated across P3–P5 as the ratio of virion-associated DelVG at passage (P_{n+1}) to intracellular DelVG at passage (P_n). **(C)** DI250 and DI250:A94G plasmids were co-transfected at a 1:1 ratio in 293 cells together with the seven remaining PR8 plasmids and a PB2 mRNA– expressing plasmid. After 48 hours, the supernatant was treated with RNase, followed by RNA extraction, DNase treatment, and RT-PCR, before amplicon sequencing. The figure shows the relative proportions of the two DelVGs incorporated into virions, following the same color scheme as in other panels. Statistical significance was determined using an unpaired t-test. ns = not significant.

To determine whether the observed replication advantage also extends to packaging, we first calculated the packaging index across passages P3–P5 and found similar ratios between DI250 and DI250:A94G across both passage windows and replicates (**Fig. 5B**). We then co-transfected both DelVG plasmids at a 1:1 ratio together with the seven remaining PR8 plasmids and a PB2 mRNA–expressing plasmid in triplicate (**Fig S5C**) and quantified the packaged DelVG sequences by Nanopore amplicon sequencing. This setup maintains a constant PB2 protein supply, eliminates competition with the full-length PB2 genome, equalizes the initial vRNA input from plasmids, and largely minimizes replication.

Before analyzing these samples, we validated our quantification approach using predefined plasmid mixtures containing both DelVGs (**Fig S5D**). Finally, sequencing of virion RNA from the 1:1 co-transfection revealed that both DIs were packaged in equal proportion, with a consistent ∼50:50 ratio across all three replicates (**Fig 5C**).

Together, these results indicate that both DelVGs are packaged with comparable efficiency when replication and WT competition are controlled, and that A94G enhances the replication and intracellular competitive advantage of DI250, accounting for its sustained dominance over time.

## Discussion

The effects of DelVGs on the fitness and phenotypes of influenza virus populations remain very poorly understood. One way to explore the interactions between WT genomes and DelVGs is to examine their co-evolutionary dynamics over time within IAV populations. Here, we used NGS to detail the complex dynamics of DelVGs during extended *in vitro* passaging under sustained high MOI conditions. By tracking individual DelVGs and WT genomes within parallel populations over 72 passages, we generated a high-resolution profile of intra-population DelVG competition. This approach, combined with *in vitro* competition assays using recombinant DelVGs, allowed us not only to identify aspects of WT/DelVG dynamics that are highly reproducible versus those that are more stochastic, but also a DelVG-specific mutation that had a profound effect on DelVG replication and competition with WT genomes.

We observed clear inter-DelVG competition primarily within the PB2 segment and, to a lesser extent, in other polymerase segments, whereas DelVG dynamics within the non-polymerase segments were more consistent with random drift. A parsimonious explanation is that deletions in polymerase segments render virions replication-deficient, enforcing strict dependence on WT complementation; under high MOI, such PB2-derived DelVGs can exploit limiting pools of the polymerase and NP, driving repeatable selective sweeps. By contrast, DelVGs in non-polymerase segments can still be replicated as RNAs in coinfected cells but typically generate semi- or non-infectious progeny (24) and face strong packaging bottlenecks that limit their potential for direct competition. This interpretation is supported by mechanistic models showing that DelVGs gain a competitive edge by sequestering limiting polymerase and NP, with the models also predicting greater competitiveness for polymerase-segment DelVGs (25, 26).

Our data indicate that A94G confers an advantage to PB2 DelVGs primarily at the replication level, which is likely contribute to its strong interfering activity. While the precise mechanism remains to be elucidated, we did not observe any bias toward in-frame deletions; however, DI250 happens to be in-frame. This raises the possibility that its encoded protein may play a role, especially given recent evidence suggesting that DelVG-derived proteins can act as dominant-negative inhibitors in cell culture (23).

Our experiments were performed in MDCK cells under controlled MOI conditions, which may not fully capture the dynamics of DelVG replication and packaging in other cell lines or *in vivo*. Moreover, our quantification relied on junction-specific qPCR of two selected DelVG clones, potentially overlooking contributions from other variants present in natural infections. While our design allowed replication to be decoupled from packaging, we cannot exclude additional factors such as RNA stability, host cell type, or segment interactions influencing the relative success of DI250:A94G over DI250. Future studies in primary human airway cells or animal models, and structural analyses of DelVG junctions, will be needed to generalize these findings and pinpoint the molecular basis of the observed differences

Prior studies have established how DelVGs shape infection by mapping deletion architectures that drive strong interference in arboviruses (ZIKV, CHIKV) and identifying DelVG-embedded features that heighten innate signaling in paramyxoviruses (27–29). Complementary evidence from plant RNA viruses shows that even single-nucleotide changes within DelVG RNAs can modulate their competitive behavior, underscoring that DelVG performance can hinge on fine-scale sequence variation and not only on deletion endpoints (30–32). Building on and integrating these insights, we show in influenza A that a SNP inside a PB2-derived DelVG repeatedly rose to high frequency over ∼72 serial high-MOI passages, enhanced DelVG competitive success, and more strongly suppressed WT replication, while the same change was lethal in the WT background with evidence of reversion pressure. To our knowledge, this is the first demonstration in influenza that a minimal point change within *a bona fide* DelVG can tune interfering capacity and expose antagonistic pleiotropy *i*.*e*. decoupled fitness optima for DelVG versus WT genomes.

Altogether, these results illuminate the co-evolutionary dynamics that govern DelVG dynamics within IAV populations and point towards actionable approaches for engineering interference-optimized DVGs for therapeutic purposes.

## Methods

### Cells and viruses

MDCK and HEK293T cells were maintained in MEM (Gibco) supplemented with GlutaMAX and 8.3% FBS (Seradigm) at 37 °C, 5% CO_2_. Recombinant A/Puerto Rico/8/1934 (PR8) was rescued in HEK293T cells using the canonical 8-plasmid influenza reverse-genetics system (33). Undiluted rescue supernatant was transferred directly onto MDCK cells, and supernatants were collected at the first appearance of cytopathic effect to produce seed stocks. Working virus stocks were then prepared by infecting MDCK cells with seed stock at an MOI of 0.0001 TCID_50_ per cell (to limit DelVG accumulation) and harvesting at 48 hpi. Supernatants were clarified at 3,000 rpm for 5 min and aliquoted (0.5 mL) for storage at −70 °C.

### Serial virus passage

Confluent MDCK were infected in replicates with PR8 at an MOI of 10 TCID50/cell, 24hpi, 500 µl of the supernatant was collected to re-infect fresh MDCK cells to generate Passage 1. We repeated this for 72 times. From the remaining supernatant of each passage, 140 µl was used for RNA extraction using QIAGEN QIAamp viral RNA mini kit according to the manufacturer’s instructions, which was kept, with the rest of the supernatant at -70°C.

### Viral cDNA amplification and sequencing

Universal RT-PCR was performed on all the samples before sequencing on Illumina MiSeq or NovaSeq using a previously described method (6). For RT-qPCR quantification of DI250, DI143, DI289 and PB2-WT gene segments, we designed and optimized specific primer/probe sets specific for each species using the IDT PrimerQuest webtool and validated efficiency and specificity using serial dilutions of plasmids encoding either species **(Fig S4 & Table 1)**. Viral RNA was extracted from cells or virions as described above and used to synthesize cDNA using the universal primer and a Verso cDNA synthesis kit (Thermo Scientific). We mixed 3µl of RNA with 8µl of H2O, 4µl of cDNA synthesis buffer, 2µl of dNTP mix (5 mM each), 1µl of universal primer (10 µM), 1µl of RT enhancer, and 1µl of Verso enzyme mix, before incubation for 50 mins at 45°C. After this, 1µl of the cDNA product was mixed with 6µl of H2O, 1µl of forward primers (18 µM), 1 µl of reverse primer (18 µM), 1 µl of specific probe (5 µM), and 10 ml of TaqMan Fast Advanced Master Mix (Thermo Scientific). The qPCR conditions used were as follows: 50°C (2 min) and 95°C (2 min), followed by 40 cycles of 95°C (1 s) and 61°C (20 s) using a qPCR QuantStudio 3 thermocycler. The qPCR in Fig 4&5 was generated using Bio-Rad CFX96 thermocycler.

**Table 1.**
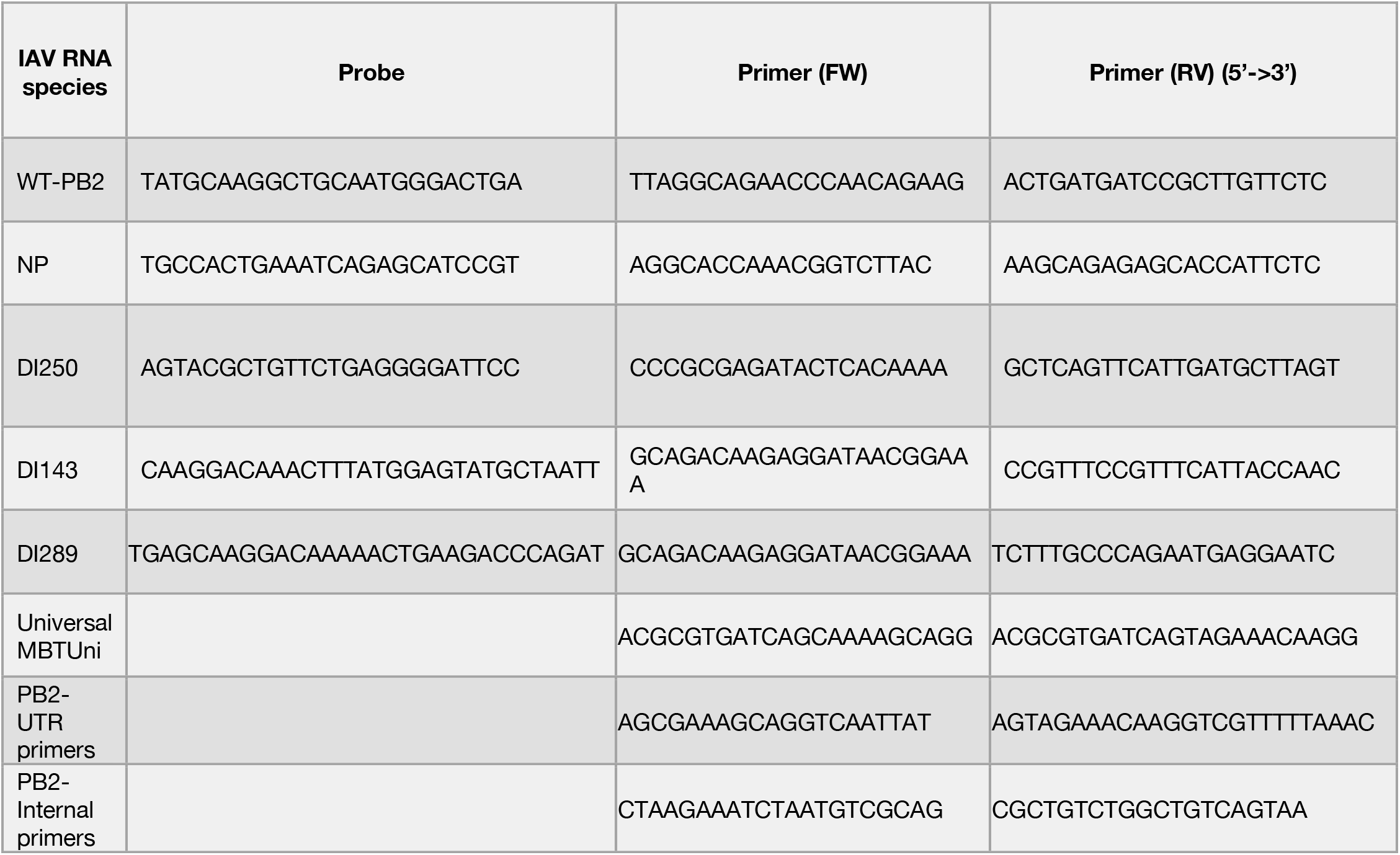
Primers and probes used in this study are listed in the 5’ to 3’ direction. The probes are labeled with 5’FAM/ZEN/3’IBFQ, and all were synthesized by Integrated DNA Technologies (IDT)

### Generation of recombinant DIPs and mutation insertion

We synthesized (Integrated DNA Technologies, Inc.) and cloned the full-length sequences of the 3 selected PB2-derived DelVGs (DI250, DI143, DI289) **(Fig 2 & S4)** into pDZ vectors and transfected them individually along with the WT-encoding versions of the remaining 7 plasmids into PB2-expressing HEK293 cells **(Fig S4)** using a standard eight-plasmid reverse-genetics approach, as previously described (6). The transfection supernatant was used to infect PB2-expressing MDCK cells (MDCK-PB2) for 48 h to generate a seed stock. After treating with DNase, we assessed the clonality of our DIP stocks by reverse transcription reactions with or without reverse transcriptase (RT), followed by a PCR amplification using specific PB2 primers **(Fig S4B & S5A)**. As expected, we found no product when the RT enzyme was left out, while distinct bands with the expected sizes appeared in the presence of RT, and no WT-derived band was observed. MDCK-PB2 cells were kindly provided by Stefan Pöhlmann and were described previously (34). For mutation insertion, the Q5 Site-Directed Mutagenesis Kit (New England Biolabs, NEB) was used.

### *In vitro* co-passaging and competition assays

To co-passage the WT and selected DelVGs **(Fig 3B)**, MDCK cells were co-infected at an MOI of 1 PB2 gene equivalent per cell for each DIP (total MOI 3) alongside 10 MOI of WT, with infections performed in triplicate. After 1 hour of adsorption at 4°C, the inoculum was removed, cells were washed, and MEM supplemented with FBS was added. Progeny virus was collected at 17–20 hpi, and a fixed volume was used to infect fresh cells. Viral RNA was extracted using the Qiagen QIAamp Viral RNA Mini Kit, and TaqMan qPCR was performed to quantify viral populations. For the competition assay **(Fig 4C, right panel)**, MDCK cells were co-infected at 0.1 MOI WT with DelVG at a 1:10 WT-to-DelVG ratio based on NP gene equivalents, and progeny was harvested at 48 hpi for plaque assay.

### Sequencing analysis of deletion junctions and data availability

Raw sequencing reads (available at #) were filtered based on quality and lengths and fed into our DI-detection pipeline for DelVG junction detection as previously described (6). Briefly, to maximize our confidence that the detected DelVGs were not derived from artifacts, we accepted only DelVGs with read support above the cutoff values, which were determined by correlation analysis between two technical replicates, i.e. independent RNA extraction of a couple of samples. Additionally, to account for variabilities between NGS datasets and segments we normalized both the NGS read and distinct junctions counts to 10^6^ mapped reads relative to the total aligned reads. Of note, throughout the analysis we carefully considered both levels of the data, the junction count and the NGS read count/support, with the former speaks more to DelVG production and the latter to DelVG replication.

To measure the WT frequency, we defined the DelVG hotspot occurrences of each segment from our NGS analysis before counting the reads that aligned outside these hotspot regions. Next, we normalized the read counts the same way we did with DelVGs to account for the differences across the NGS datasets. We validated this method by using simulated NGS datasets from our previous study that were derived from Cal07 and contained a defined proportion of DelVG and WT reads (6), and found no significant difference between the expected and observed (Chi Square = 0.42) in three different libraries contain different amount of DelVG junctions **(Fig S6)**. To profile mutations in DelVGs, we leveraged their short length and junction placement, which are typically fully covered by ∼250-nt reads. Using DelVG sequences as references, we extracted reads spanning the deletion junction and required ≥100 nt of coverage on both sides before conducting variant analysis.

Finally, the mutations were analyzed using `LowFreq` (35) and `SNPGenie` (36), and then visualized in circular plots using circle `packcircles` and `ggplot2` packages in R.

### Shannon Diversity

The probability of each DelVG was calculated by dividing the NGS read support of each DleVG by the total DelVG reads, before applying the following equation to calculate the diversity of each population. Shannon Diversity takes into consideration both the NGS and junction counts; the higher the value the higher the diversity.

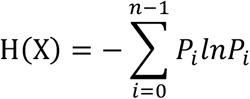

### Minigenome replicon

HEK293T cells were seeded at 1.5 × 10^5^ cells per well in 24-well plates. After 24 hours, the cells were co-transfected using PolyPlus JetPrime with plasmids encoding the influenza PR8 replication machinery proteins, along with pCI-NP and two additional plasmids: pPolI-Firefly, which expresses the Firefly luciferase coding sequence in negative polarity flanked by the 5′ and 3′ non-coding regions of the NS segment, and pTK-Renilla, driven by a cellular promoter. All plasmids were generously provided by Nadia Naffakh and have been previously described (37). Six hours post-transfection, the medium was replaced with fresh medium containing 10% FBS and 1% Pen/Strep. After 24 hours of incubation at 37°C, luciferase activity was measured using the Renilla Luciferase Assay System (Promega) and a GloMax luminometer.

### The hemagglutination (HA) assay

The HA assay was performed in a round-bottom 96-well plate. Serial two-fold dilutions of the virus samples were prepared in PBS (+/+) to a final volume of 50 μL per well, including virus-free negative controls. Turkey red blood cells (RBCs) were washed by resuspending them in 15 mL of PBS (+/+) in a 15 mL conical tube, followed by centrifugation at 1,000 × g for 5 minutes. The supernatant was discarded, and the RBCs were resuspended to a final concentration of 1%. Next, 50 μL of the 1% RBC suspension was added to each well, and the plates were incubated at 4°C for 30–60 minutes. The endpoint dilution showing complete hemagglutination was recorded.

### DI nomenclature

In this study we follow the nomenclature introduced by Dimmock and Easton (14), where defective interfering (DI) RNAs are designated by the segment of origin and the length of the sequence retained from the 3′ end of the positive-sense RNA before the internal deletion. For example, the prototype DI244 (also written as DI 1/244) derives from segment 1 (PB2) and retains the first 244 nucleotides from the 3′ end of the positive strand, together with the terminal nucleotides from the 5′ end, thereby defining the breakpoints of the internal deletion.

## Acknowledgements

This work was generously supported by the Defence Advanced Research Projects Agency under contract DARPA-16-35-INTERCEPT-FP-018, the National Institutes of Health under grants R01AI139246, R01AI179910, and 1U01AI186993, and The Agency for Science, Technology and Research (A*STAR), Infectious diseases labs (IDL) in Singapore.

## Supplementary figures

**Fig S1.**
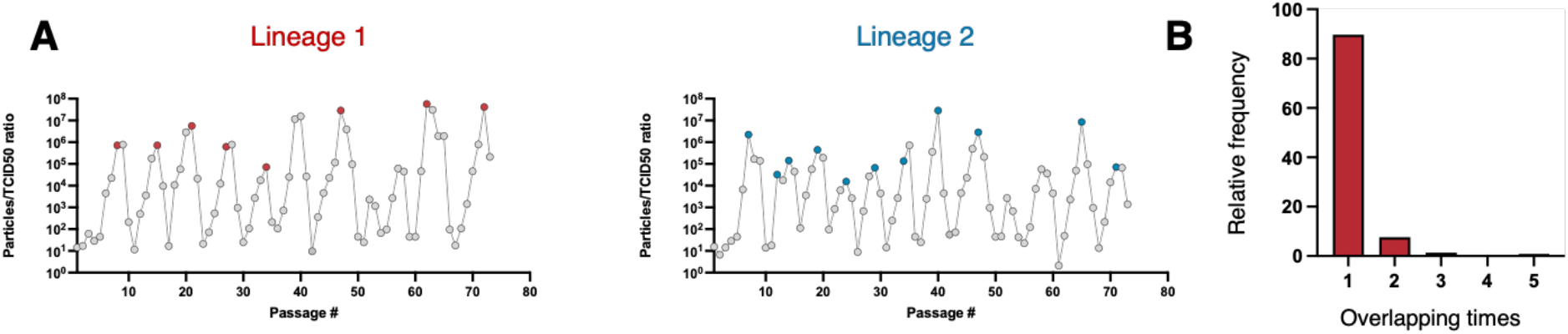
DelVG Junction analysis. (A) Plots showing the particles to infectivity ratio at each passage in the long-passaging experiment with the samples selected for NGS sequencing are highlighted in both lineages. (B) The percentage frequency of the overlap occurrences in PB2 between the 6 replicates relative to the total overlap times, 1 = unique junction, 2 = found in 2 replicates, 3 = found in 3 replicates, and so on.

**Fig S2.**
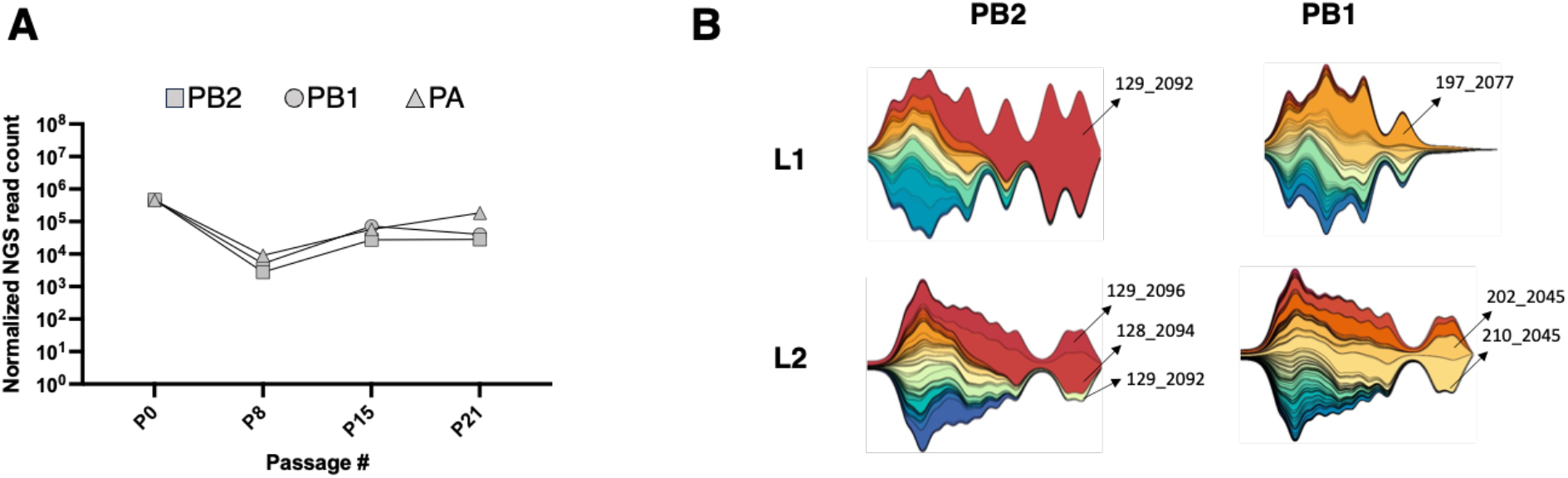
WT read kinetics and mapping of dominant DelVG junctions. (A) The normalized read counts of WT-derived NGS reads for segments 1-3 at the first 4 DIP-enriched passages (B) The same streamplots in Fig 3 with the junction sites of the highest competitive ones are indicated.

**Fig S3.**
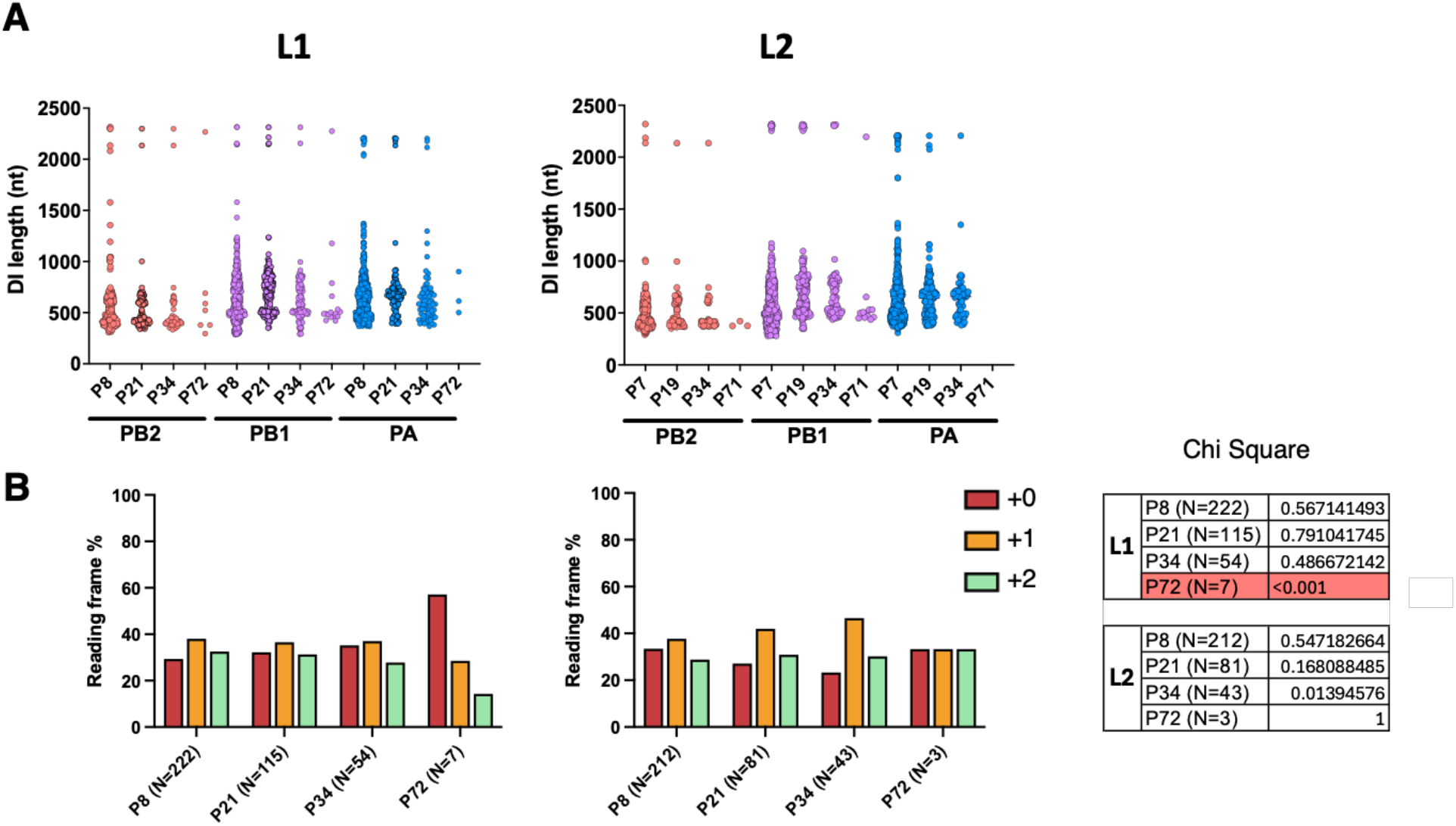
The analysis of DelVG size and reading frame in both lineages. (A) DelVG size analysis, where the size of the junctions was calculated across four representative passages for the polymerase segments and plotted alongside each other. (B) A bar chart displaying the frequency (%) of all possible reading frames for all DelVGs across the four passages. The “+0” denotes the reading frame identical to the parental genome, while “+1” and “+2” indicate shifts in the reading frame by one or two nucleotides, respectively.

**Fig S4.**
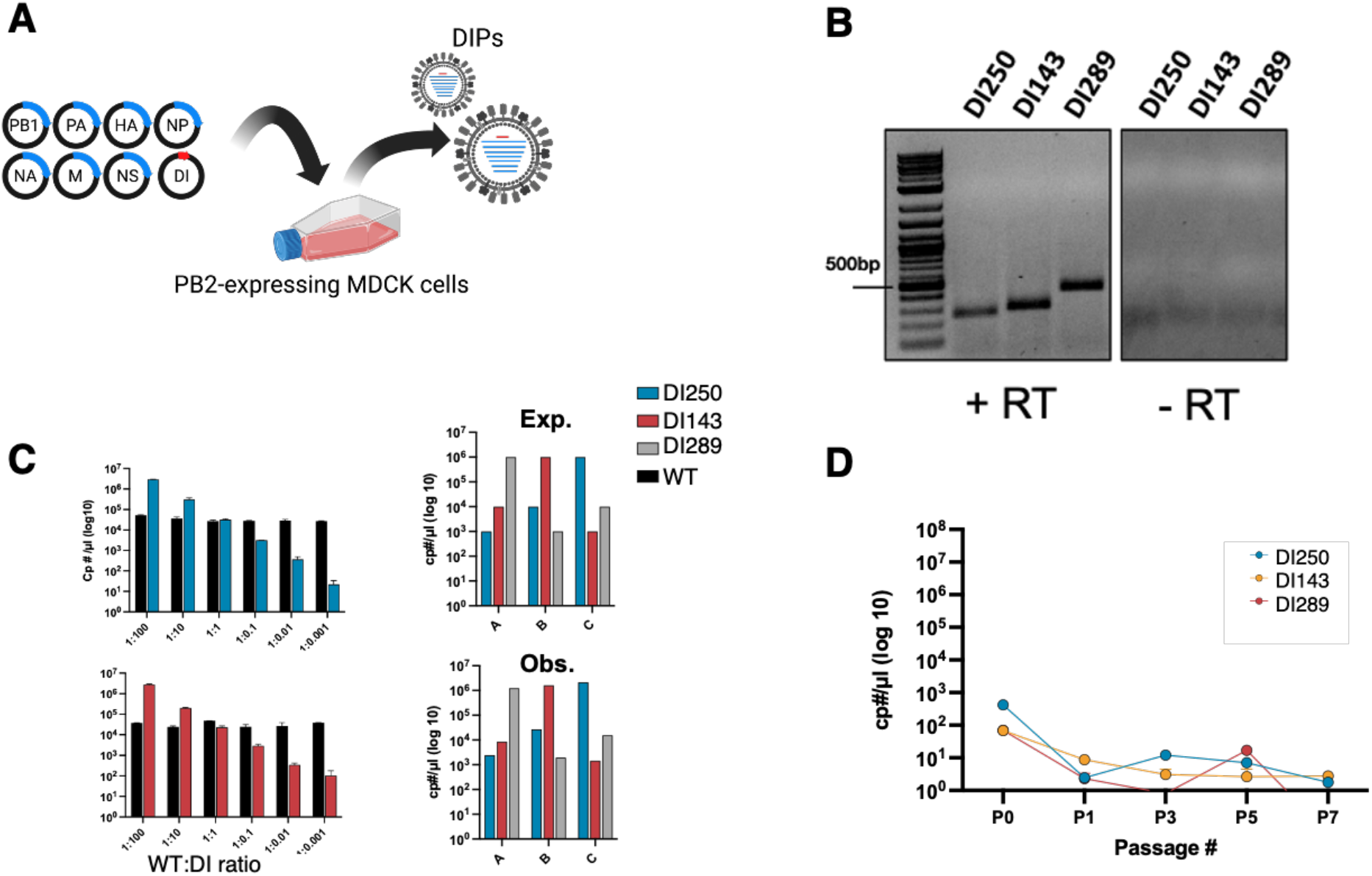
The selection of DelVGs for clonal DIPs preparation and the experimental verifications. (A) A cartoon drawing illustrates the experimental setup for DIP generation. (B) PCR amplification gel analysis of the DIPs resulted from B. DIPs RNA was extracted, followed by DNase treatment and cDNA synthesis (with or without the reverse transcriptase) and PCR amplification using specific internal PB2 primers (see the primers table). (C) Left panel displays the optimization of primers and probes targeting specific DelVGs selected for the study, showing the detected copy numbers obtained from defined mixtures of WT and DelVG plasmids. The data presented correspond to DI250 and DI289. Right panel: Plasmids from three selected DelVGs were combined in varying predefined ratios (Exp = expected) and quantified using qPCR (Obs = observed) (D) Negative control showing the inability of DIP to grow in the absence of WT

**Fig S5.**
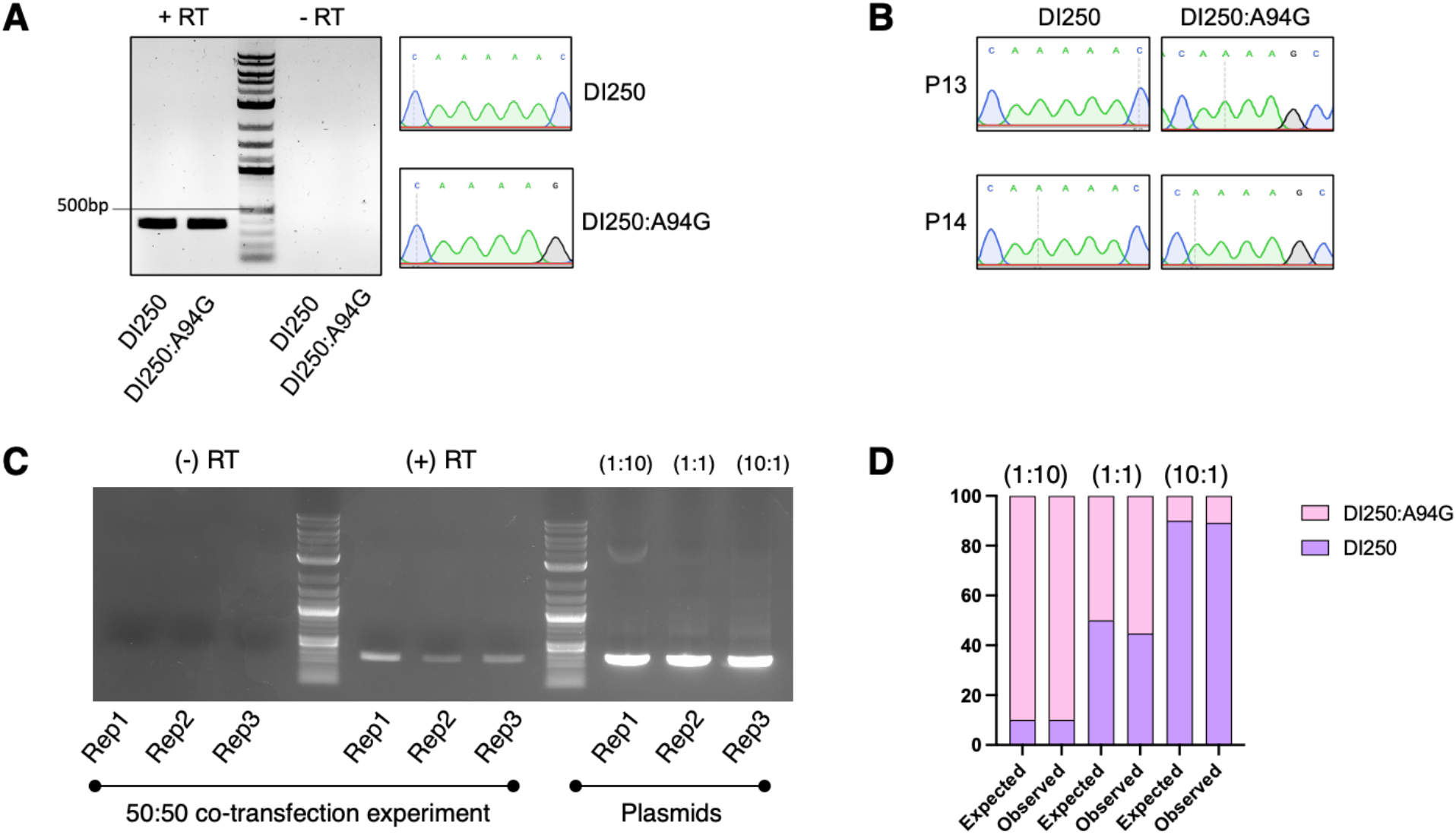
Validation and quantification of A94G by Sanger and Nanopore sequencing. (A) The same method in Fig S4 was used with the specific PB2-UTR primers (see primers table), and sanger sequencing was employed to confirm the A94G mutation. (B) Sanger sequencing of DI250 and DI250:A94G at passages 13 and 14 in Rep 1(Fig. 4D) was performed using junction-specific primers, and the same results were obtained in Rep 2. (C) DI250 and DI250:A94G plasmids were co-transfected at a 1:1 ratio with the seven remaining PR8 plasmids and a PB2 mRNA-expressing plasmid. After 48 hours, the supernatant was RNase-treated to remove unprotected RNA, followed by RNA extraction and DNase treatment before RT-PCR. The resulting bands from the co-transfection and the predefined mixed plasmids were visualized on a gel, excised, and sent for Nanopore amplicon sequencing. (D) Sequencing analysis of the plasmid mixes from panel C shows that the observed ratios closely align with the expected ones.

**Fig S6.**
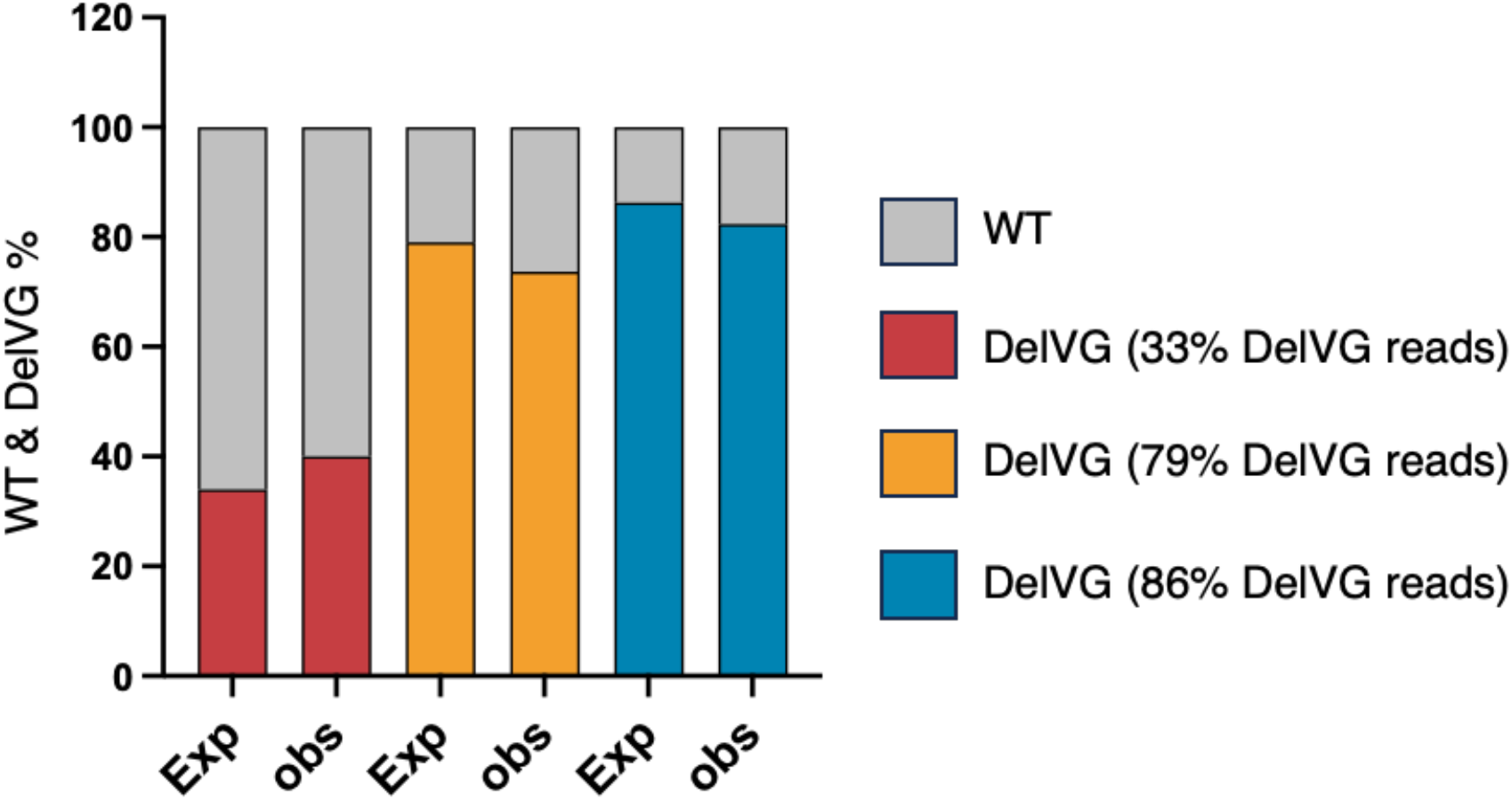
Data simulation control. Simulated datasets with known WT:DI ratios were used to validate counting the WT-derived NGS read count. The expected ratio of DelVG reads is indicated) The staked bar is showing the WT:DI NGS read ratio between the expected (defined in the datasets) and observed (resulted from analysing the datasets).

